# AdmixSim: A Forward-Time Simulator for Various and Complex Scenarios of Population Admixture

**DOI:** 10.1101/037135

**Authors:** Xiong Yang, Xumin Ni, Ying Zhou, Wei Guo, Kai Yuan, Shuhua Xu

## Abstract

**Background:** Population admixture has been a common phenomenon in human, animals and plants, and plays a very important role in shaping individual genetic architecture and population genetic diversity. Inference of population admixture, however, is challenging and typically relies on *in silico* simulation. We are aware of the lack of a computer tool for such a purpose, especially a simulator is not available for generating data under various and complex admixture scenarios.

**Results:** Here we developed a forward-time simulator (AdmixSim) under standard Wright Fisher model, which can simulate admixed populations with: 1) multiple ancestral populations; 2) multiple waves of admixture events; 3) fluctuating population size; and 4) fluctuating admixture proportions. Results of analysis of the simulated data by AdmixSim show that our simulator can fast and accurately generate data resemble real one. We included in AdmixSim all possible parameters that allow users to modify and simulate any kinds of admixture scenarios easily so that it is very flexible. AdmixSim records recombination break points and trace of each chromosomal segment from different ancestral populations, with which users can easily do further analysis and comparative studies with empirical data.

**Conclusions:** AdmixSim is expected to facilitate the study of population admixture by providing a simulation framework with flexible implementation of various admixture models and parameters.

## Background

Human migration resulted in population differentiation, and many populations, especially those living in different continents, have been isolated for quite a long time. However, subsequent migrations that occurred over the past millennia have resulted in gene flow between previously separated human subpopulations. This has been a common phenomenon throughout the history of modern humans, as previously isolated populations often come into contact through colonization and migration. As a result, admixed populations came into being when previously mutually isolated populations met and inter-married. In human, for instance, it was estimated there are over 100 admixture events occurred in recent 4000 years [1], which plays an important role in shaping genetic diversity of modern humans. Therefore, study of population admixture will shed light on human genetic history and has many implications in medical research. However, inferring population admixture relies strongly on simulation. *In silico* simulation is useful in testing population genetic model, studying recombination, linkage disequilibrium, ancestry tracking, and admixture dynamics in admixed populations. A bunch of forward- and backward-time simulators [2–7] has been developed in recent years. Some of the backward-time simulator (most of them are coalescent based), for example *ms*[2], could simulate population admixture under simple scenarios. However, it’s very complex or even impossible to simulate an admixed population with fluctuating population size(s) or fluctuating gene flow(s) generation by generation. Some of the forward-time simulator, for example *SFS_CODE*[7], could also simulate admixture under simple scenarios, but they also suffered from the same problems of coalescent based simulators. To our knowledge, there is no simulator that focuses on simulating and tracking the dynamics of recombination and ancestry in admixed populations, and allowing change population size(s) and gene flow(s) generation by generation.

## Implementation

### A Generalized Admixture Model

Population admixture is accomplished by receiving gene flow(s) from ancestral populations either continuously or discontinuously. To make the modeling of population admixture more general, we can model this process generation by generation, in which, if the admixed population does not receive further gene flow(s) in a particular generation, we set the strength of gene flow(s) to 0. Given an admixed population with *K*(*K* > 1) ancestral populations and was formed *T* (*T* > 0) generations ago can be well modeled by a *K×T* matrix *M*. The *m_ij_* in *M* denotes the strength of gene flow from *jth* ancestral population at *ith* generation, where *m_ij_*fulfills two requirements: 1) 0 ≤ *m_ij_*≤ 1; and 2) ∑ *m_ij_*= 1 when *i* = 1.

### Simulation Process

In a recent admixed population, novel mutations and selections have negligible impacts on shaping the genetic diversity on whole genome scale, here we neglected mutations and selections in our simulation. Recombination is modeled as Poison process along chromosome (chromosomal end is ignored) with rate 1 (unit in Morgan). In a certain generation, *i*, the population size of admixed population is *N*, and the rate of gene flow from *jth* ancestral population is *m_ij_*, we generate individuals in current generation by two steps:

1. For gene flow from the *jth* ancestral population, we randomly sample *N×m_ij_* individuals from the *jth* ancestral population, and repeat this procedure for all the gene flow events at current generation and the rest of individuals are randomly sampled from the admixed population in previous generation;
2. With the sample pool generated in step 1), we randomly choose two individuals in the pool, then randomly choose one of the chromosomes from one individual, pair and recombine it with the one randomly chosen in another individual, to form a new chromosome pair. Repeat these procedures until *N* individuals are generated.

Individuals are generated by the two steps generation by generation. At the end of simulation, n individuals are randomly sampled; the start, end positions and the ancestry of a chromosome segment from which are recorded. In addition, the haplotypes for each individual are also recorded for further studies.

## Results and Discussion

As previous study has already deduced the theoretical distribution of the length of ancestral chromosomal segment (LACS) under HI model [8], it is straightforward to test the performance of our simulator. Under HI model, the distribution of LACS from ancestral population with ancestry contribution *m* is *(1-m)Te^-(1-m)Tx^*. Here we simulated an admixed population with constant *Ne*=5000, admixed 100 generations ago, following HI model, in which ancestral population 1 contributes 25% of the total ancestry. As expected, the distribution of LACS simulated (dots) matches the theoretical distribution of LACS (dashed line) well for ancestries from both ancestral population 1 and population 2 (Figure 1).

**Figure 1.**
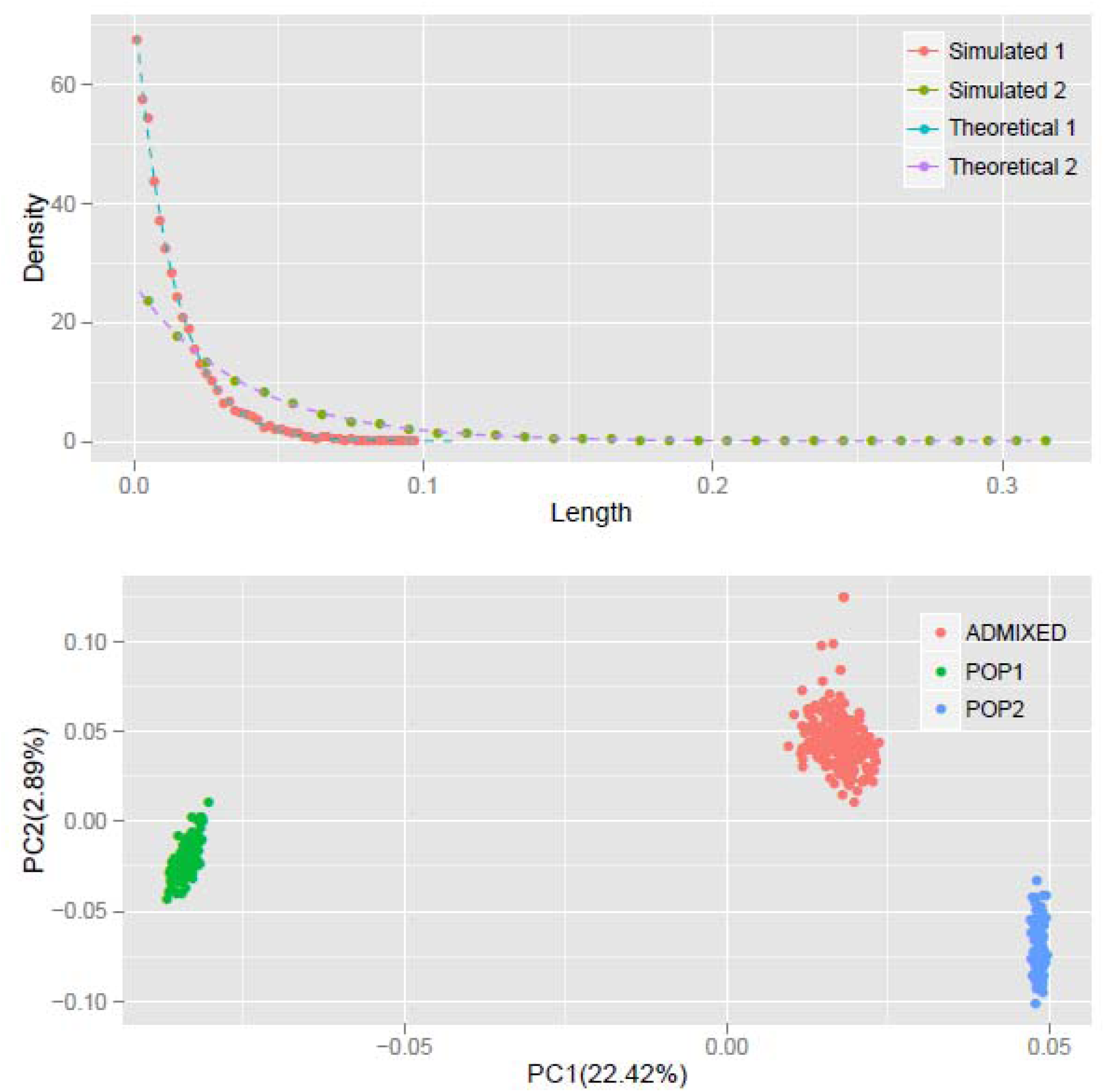
The distribution of LACS and PCA of the simulated data. Top: red and green dots denote the distribution of LACS from ancestral population 1 and ancestral population 2; blue and purple dashed lines are the theoretical distributions of LACS derived from *(1-m)Te^-(1-m)Tx^*, where *m* is the ancestry contribution of that ancestral population and *T* is the generation since admixture. Bottom: green and blue dots denote the individuals from ancestral populations and the red dots denote individuals in admixed population.

Principal component analysis (PCA) with *smartpca*[9, 10] shows that first principal component (PC) separates ancestral population 1 and ancestral population 2, just as expected and usually observed in real data. Simulated individuals from the two ancestral populations cluster within their own groups, and the admixed individuals cluster between two ancestral populations along PC1 (Figure 1). As ancestral population 2 contributes more to the admixed population, the distance between individual from admixed population and individual from ancestral population 2 is much closer than those from ancestral population 1 to the admixed individuals. All these results observed in simulated data resemble what observed in real data, which indicates our simulator does generate correct dataset which resembles the real one.

Moreover, in the result of *ADMIXTURE* analysis [11] of the simulated data, we can clearly observe the assignment of ancestries of the admixed individuals: blue represents ancestry from ancestral population 1 and green represents ancestry from ancestral population 2. The ancestry contributions from ancestral population 1 vary from 18% to 30% (Figure S1), which also resembles what observed in real data.

**Figure S1.**
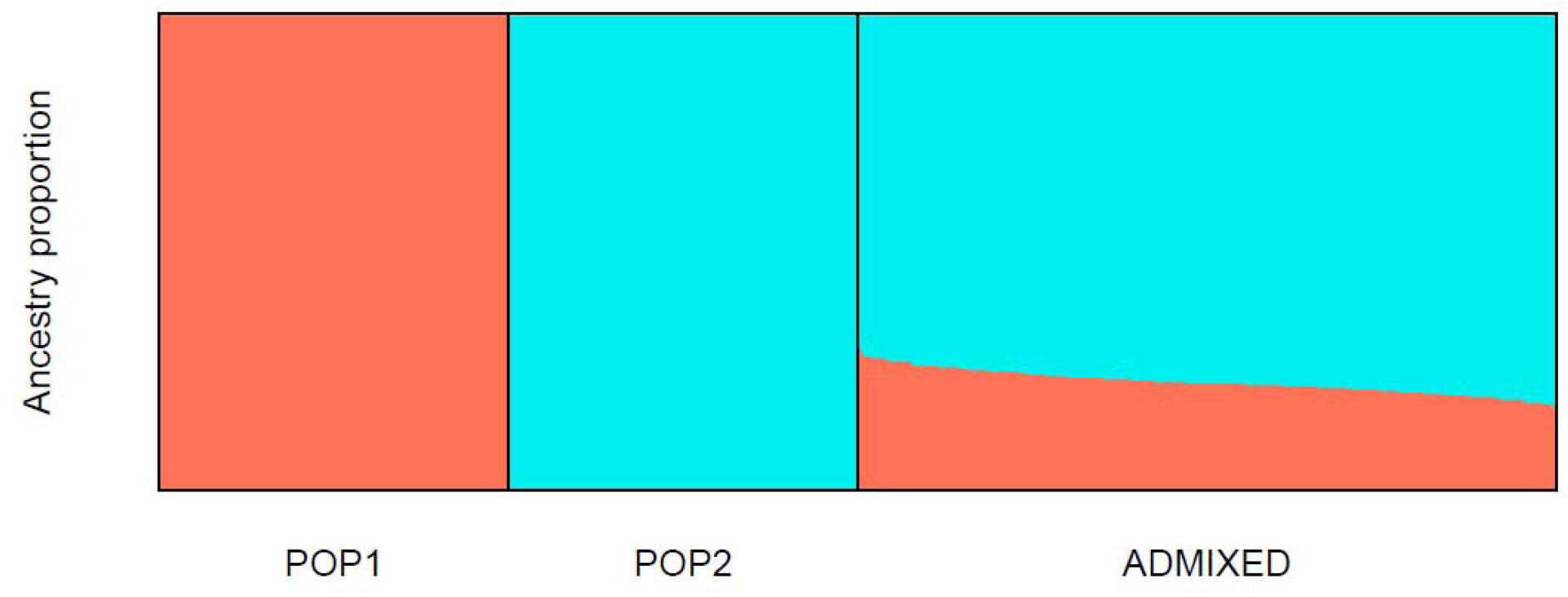
*ADMIXTURE* analysis of simulated data. Blue and green denote two ancestral components and the admixed individuals show combination of the ancestries from two ancestral populations. Tutorials for AdmixSim

The time complexity of AdmixSim is *O(L), O(N)* and *O(T^2^)*, thus in theory, the running time of AdmixSim is linearly decided by the chromosome length (L) and the population size (N) simulated; and quadratic depends on the generations since admixture (T). Benchmark tests are carried on a laptop with *Intel Core^TM^Duo* CPU @ 2.0GHz, 2 Gb RAM and Ubuntu 12.04 32-bit operation system. Running time is recorded by Linux command *time* and the *user* time is collected and compared. Each time, only one parameter is allowed to be variable. For example, to test the impact of generation since admixture on running time, we run a series of tests in which generation ranges from start to end, increased by 1 step size each time. The details of parameters setting for each test can be found in Table 1. The results show that the AdmixSim runs very fast even with large population size, and the running time shows a trend as expected (Figure 2).

**Table 1.** Detailed parameters of benchmark test. First column is the parameter to be tested; second and the following two columns are the ranges of the parameters (start, end and increasing step); last column is all other parameters setting to constants. N is population size, g is generation since admixture, k is the number of ancestral populations, l is the chromosome length (unit in Morgan), and n is the number of individuals sampled from admixed population at the end of simulation.

| Parameter | Start | End | Step | Other fixed settings |
| --- | --- | --- | --- | --- |
| generation (g) | 5 | 150 | 5 | N=5000, k=2, l=1 and n=200 |
| chromosome length (l) | 1 | 30 | 1 | N=5000, k=2, g=20 and<br>n=200 |
| population size(N) | 500 | 20000 | 500 | k=2, l=1, g=20 and n=200 |

**Figure 2.**
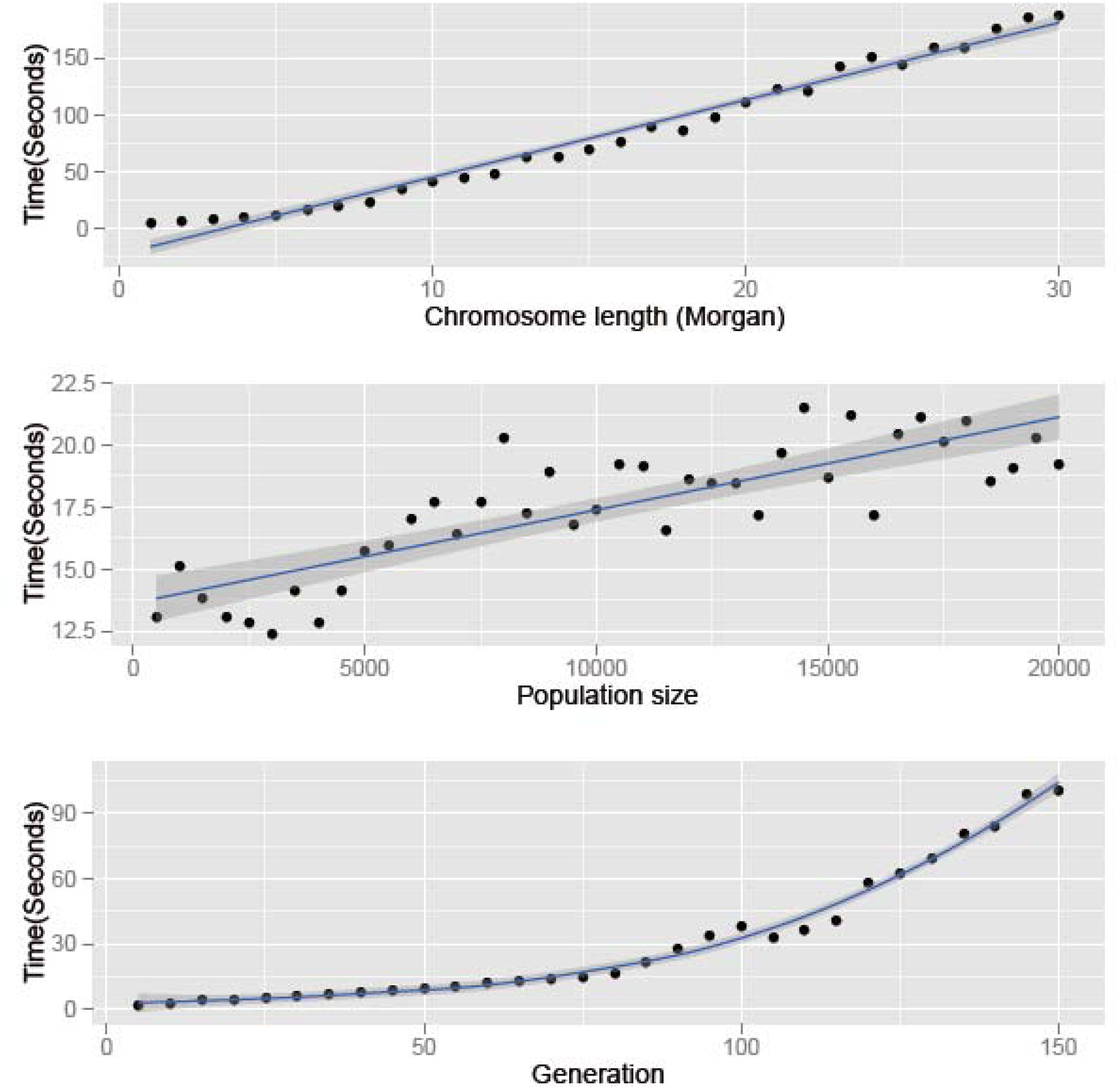
Running time of AdmixSim. Top: running time versus chromosome length simulated; middle: running time versus population size simulated; and bottom: running time versus time in generation since admixture.

## Conclusions

Here we developed a fast and flexible simulator aiming for modeling various and complex scenarios of population admixture, with which can simulate an admixed population with: 1) multiple ancestral populations; 2) multiple waves of admixture events; 3) fluctuating population sizes; and 4) fluctuating admixture proportions. With AdmixSim, the users can not only easily simulate an admixed population under the three typical admixture models, i.e. hybrid isolation (HI) model, gradual admixture (GA) model, and continuous gene flow (CGF) model as described in previous studies [12], but also simulate an admixed population with more complex scenarios, for example, three-way admixture with multiple waves of gene flows. This simulator is expected to facilitate study of population admixture thus can greatly help us to understand the processes of human migration, admixture and evolution, which further gives insights into both evolutionary and medical studies of human genetic diversity.

## Availability and requirements

Project name: AdmixSim

Project home page: http://www.picb.ac.cn/PGG/resource.php

Operating system(s): Platform independent

Programming language: Java Other requirements: Java 1.6 or higher.

License: GNU General Public License V3Any restrictions to use by non-academics: license needed

## Abbreviations

CGF: Continuous gene flow
GA: Gradual admixture
HI: Hybrid isolation
LACS: Length of ancestral chromosomal segment
PCA: Principal component analysis
PC: Principal component

## Competing interests

The authors declare that they have no competing interests.

## Author contributions

SX conceived the study. XY designed and implemented the AdmixSim with contribution from XN WG and YZ. XY, SX wrote the manuscript. XY, XN and WG analyzed the time complexity and the performance of the simulator with contribution from KY. All authors read and approved the final manuscript.

## Supplementary Materials

Supplementary materials include Figure S1 and Text S1.

## Acknowledgements

These studies were supported by the National Science Foundation of China (NSFC) grants (91331204; 31171218), by the Strategic Priority Research Program of the Chinese Academy of Sciences (CAS) (XDB13040100). This research was supported in part by the Ministry of Science and Technology (MoST) International Cooperation Base of China and by National Center for Mathematics and Interdisciplinary Sciences (NCMIS), Academy of Mathematics and Systems Science, CAS. W.G was supported by the Fundamental Research Funds for the Central Universities (2011JBZ019). S.X. is Max-Planck Independent Research Group Leader and member of CAS Youth Innovation Promotion Association. S.X. also gratefully acknowledges the support of the National Program for Top-notch Young Innovative Talents of The “*Wanren Jihua*” Project.

